# The aphelid-like phagotrophic origins of fungi

**DOI:** 10.1101/233882

**Authors:** Guifré Torruella, Xavier Grau-Bové, David Moreira, Sergey A. Karpov, John A. Burns, Arnau Sebé-Pedrós, Eckhard Völcker, Purificación López-García

## Abstract

Aphelids are poorly known phagotrophic parasites of algae whose life cycle and morphology resemble those of the widely diverse parasitic rozellids (Cryptomycota, Rozellomycota). In previous phylogenetic analyses of RNA polymerase and rRNA genes, aphelids and rozellids formed a monophyletic group together with the extremely reduced parasitic Microsporidia, named Opisthosporidia, which was sister to Fungi. However, the statistical support for that group was always moderate. We generated the first transcriptome data for one aphelid species, *Paraphelidium tribonemae*. In-depth multi-gene phylogenomic analyses using various protein datasets place aphelids as the closest relatives of Fungi to the exclusion of rozellids and Microsporidia. In contrast with the comparatively reduced *Rozella allomycis* genome, we infer a rich, free-living-like aphelid proteome, including cellulases likely involved in algal cell-wall penetration, enzymes involved in chitin biosynthesis and several metabolic pathways. Our results suggest that Fungi evolved from a complex aphelid-like ancestor that lost phagotrophy and became osmotrophic.

---

Aphelids constitute a group of diverse, yet poorly known, parasites of algae^1,2^. Their life cycle and morphology resemble those of zoosporic fungi (chytrids) and rozellids (Cryptomycota/Rozellosporidia), another specious group of parasites of fungi and oomycetes^3,4^. Unlike fungi, which are osmotrophs, aphelids and rozellids are phagotrophs, feeding on the host’s cytoplasm. Combined RNA polymerase and rRNA gene trees^5^ suggested that aphelids and rozellids relate to Microsporidia, extremely reduced parasites with remnant mitochondria^6^. Accordingly, aphelids, rozellids and Microsporidia were proposed to form a monophyletic clade sister to Fungi, called Opisthosporidia^1^. Microsporidia would have subsequently lost the ancestral opisthosporidian phagotrophy. However, the limited phylogenetic signal of those genes combined with microsporidian fast-evolving sequences have resulted in incongruent tree topologies, showing either rozellids^5,7^ or aphelids^8^ as the earliest-branching lineages of Opisthosporidia. Furthermore, the support for the monophyly of Opisthosporidia was always moderate, leaving unresolved the relative order of emergence of the recently recognized and highly diverse lineages of eukaryotes that are related to fungi. To improve the phylogenetic signal for an accurate placement of aphelids in the opisthokont branch containing fungi, nucleariids, rozellids and Microsporidia (usually referred to as Holomycota), we have generated the first transcriptome data for one aphelid species, *Paraphelidium tribonemae*^2^. Phylogenomic analysis including, in particular, data from the *Rozella allomycis* genome^9^, should allow to validate or refute the monophyly of Opisthosporidia and determine their relative order of emergence along this line.

## Results

### Aphelids occupy a deep pivotal position and are sister to Fungi

To generate the aphelid transcriptome and because *P. tribonemae* has a complex life cycle (Fig. 1), we maximized transcript recovery by constructing two cDNA libraries corresponding to young and old enrichment cultures of its host, the yellow-green alga *Tribonema gayanum*, infected with the aphelid. Accordingly, transcripts corresponding to zoospores, infective cysts, trophonts, plasmodia, sporangia and sporocysts were represented^2^. After paired-end Illumina HiSeq2500 sequencing, we assembled a metatranscriptome of 68,130 contigs corresponding to the aphelid, its host and bacterial contaminants. After a sequential process of supervised cleaning, including the comparison with a newly generated transcriptome of the algal host, we obtained a final dataset of 10,439 protein sequences that were considered free of contaminants. The final predicted proteome (*Paraphelidium tribonemae* version 1.5; Supplementary Table 1) was 91.4% complete according to BUSCO^10^. We found no stop codons interrupting coding sequences. Therefore, in contrast to *Amoeboaphelidium protococcarum*, for which the TAG and TAA stop triplets appear to encode glutamine^5^, *P. tribonemae* has the canonical genetic code.

**Figure 1.**
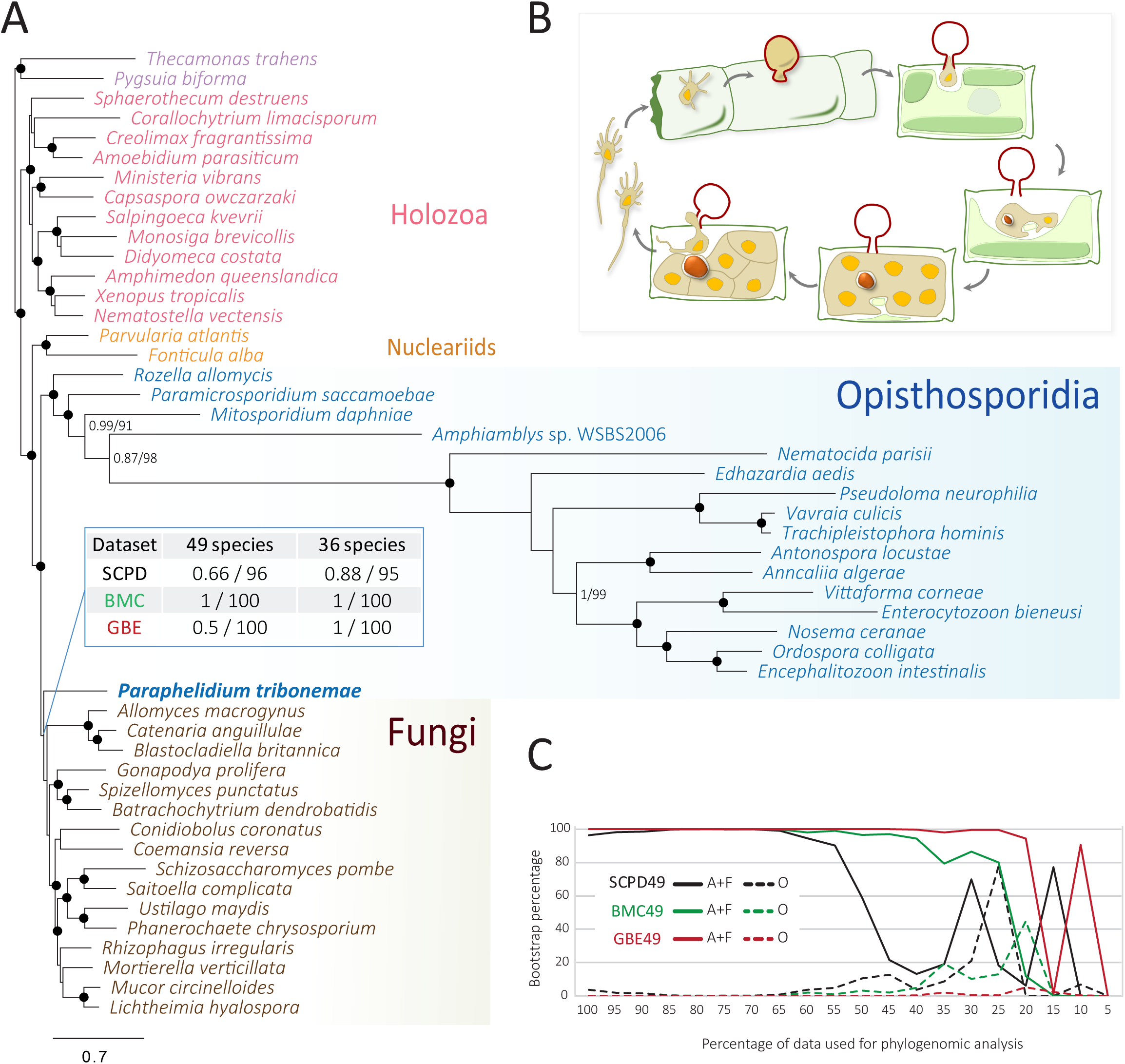
Phylogenomic analyses and cell cycle of the aphelid *Paraphelidium tribonemae*. (A) Bayesian phylogenetic tree based on single-copy protein domains for 49 species (SCPD49) inferred using a CAT-Poisson model. Statistical supports indicated at some crucial nodes correspond, from left to right, to PhyloBayes Bayesian posterior probabilities andIQ-TREE ML ultrafast-bootstrap support using the C60 model. Branches with maximum support values (BPP = 1 and UFBS = 100%) are indicated by black circles. The support for the monophyly of *Paraphelidium* and Fungi using other datasets (BMC^18^, GBE^8^) and removing the fastest-evolving microsporidian sequences (36 instead of 49 species) is shown at the corresponding node. (B) Schematic cell cycle of *P. tribonemae*. Briefly, infecting cysts (red wall), deliver an amoeboid trophont to an algal filament cell via an infection tube; the trophont engulfs the algal cytoplasm by phagocytosis, leaving a growing residual body (dark red particle); after nuclear and cell division, a mature sporangium releases amoeboflagellated zoospores (occasionally amoeboid only) that get out the algal cell wall and close the life cycle^2^. (C) Evolution of IQ-TREE ML UFBS support for the sisterhood of aphelids and Fungi (A+F) and the monophyly of Opisthosporidia (O) as a function of the proportion of fast-evolving sites removed from the dataset. All the phylogenomic trees can be seen in Supplementary Fig. 1.

We incorporated *P. tribonemae* orthologs to a dataset of 93 single-copy protein domains (SCPD dataset) previously used for phylogenomic analysis of basal-branching opisthokonts^11^, updating it with several microsporidian species including the early-branching *Mitosporidium daphniae*^12^, the metchnikovellid *Amphiamblys* sp.^8^ and the recently released genome of *Paramicrosporidium saccamoeba*^17^. We then generated Bayesian inference (BI) trees reconstructed under the CAT-Poisson mixture model^13^ and Maximum Likelihood (ML) trees reconstructed under C60, the closest mixture model to CAT^14^, for a taxon sampling including 49 species (22,976 positions). The two phylogenomic analyses yielded the same topology. We recovered the monophyly of all opisthokont lineages previously reported^11,15^. In contrast with previous analyses based on 18S rRNA and RNA polymerase genes^5,7^, Opisthosporidia appeared as paraphyletic and the aphelid was placed as sister to Fungi (Fig. 1). However, neither the Bayesian posterior probability (0.66) nor the ML bootstrap support (ultrafast bootstrapping 96%) was maximal. To limit potential long-branch attraction artefacts derived from the inclusion of fast-evolving Microsporidia, we built a reduced dataset without the thirteen fastest-evolving microsporidian species (SCPD36, 36 species; 24,413 positions). Bayesian and ML trees again reproduced the same topology (Supplementary Figs. 1c and d) with equally moderate support values (Fig. 1).

To try resolving the phylogenetic position of the aphelid with higher support, we analyzed two additional datasets previously used for phylogenomic analyses of Microsporidia with the same two sets of 49 and 36 opisthokont species. The resulting datasets were called BMC49 and BMC36 for a set of 53 protein markers^18^ which, after concatenation, yielded 15,744 and 16,840 amino acid positions, respectively; and GBE49 and GBE36 for datasets with 259 protein markers^8^, which resulted in 87,045 and 96,029 amino acids, respectively. For all datasets, Bayesian and ML trees recovered the sisterhood of *Paraphelidium* and Fungi (Supplementary Fig. 1) and, for the two datasets without fast-evolving Microsporidia, posterior probabilities and ML bootstrap support were maximal (Fig. 1, Supplementary Table 2). In addition, we carried out alternative topology tests for the three datasets containing 49 species. All of them supported the aphelid sisterhood to Fungi (Supplementary Table 2). Finally, to minimize systematic error in ML analyses due to the inclusion of fast-evolving sites in our protein datasets, we progressively removed the 5% fastest-evolving sites and reconstructed the phylogenomic trees. The support for the monophyly of Fungi and the aphelid was very high until 50% (SCPD dataset) or up to 80% (GBE dataset) of sites were removed (Fig. 1c).

Collectively, our phylogenomic analyses support the sisterhood of aphelids and Fungi, although the branch joining the two groups is very short. Opisthosporidia, united by their ancestral phagotrophy, appear as paraphyletic.

### Enzymes involved in cell-wall synthesis and degradation

Despite secondary losses in some fungi, the presence of chitin in cell walls was for long considered a typical fungal trait^19^. However, chitin is also present in the cell wall of many other protists across the eukaryotic tree, implying that the machinery for chitin synthesis/remodeling originated prior to the radiation of Fungi and other eukaryotic lineages^19^. Microsporidia and *Rozella* are also able to synthesize chitin^9,20^ but, unlike fungi, which possess chitin cell walls during the vegetative stage, they produce chitin only in cysts or resting spores^21^. Staining with fluorescently-labeled Wheat Germ Agglutinin (WGA) showed that *Paraphelidium* also possesses chitin in the wall of infecting cysts, but not in zoospores or, unlike fungi, trophonts (Fig. 2a-b). In agreement with this observation, we identified homologs of chitin synthases, chitin deacetylases, chitinases and 1,3-beta glucan-synthases in the *Paraphelidum* transcriptome (Supplementary Fig. 2a-d). Specifically, we detected seven homologous sequences (including all alternative transcripts such alleles or splice variants) of division II chitin synthases^11,20,22,23^ in *Paraphelidium* corresponding to at least six distinct peptides (*Rozella* contains only four^9^). Three of them clustered with class IV chitin synthases, including Microsporidia and *Rozella* homologs (Supplementary Fig. 2a). The remaining four sequences branched within class V/VII enzymes^23^, two of them (probably corresponding to a single polypeptide) forming a deep-branching group with fungal, mostly chytrid, sequences (Fig. 2a). Class V enzymes include a myosin motor thought to intervene in polarized fungal hyphal growth that has been hypothesized to take part in the formation of the germ tube in aphelids and rozellids^12^. Class V chitin synthases were lost in Microsporidia (with the exception of *Mitosporidium*, still retaining, like *Rozella*, one homolog), endowed instead with highly specialized polar tube extrusion mechanisms^12^. Neither spore wall nor polar tube proteins specific to Microsporidia^24^ occurred in the *Paraphelidium* transcriptome. Therefore, our data (Supplementary Fig. 2a; Supplementary Tables 1 and 5) lend credit to the hypothesis that class V chitin synthases are involved in germ tube polar growth.

**Figure 2.**
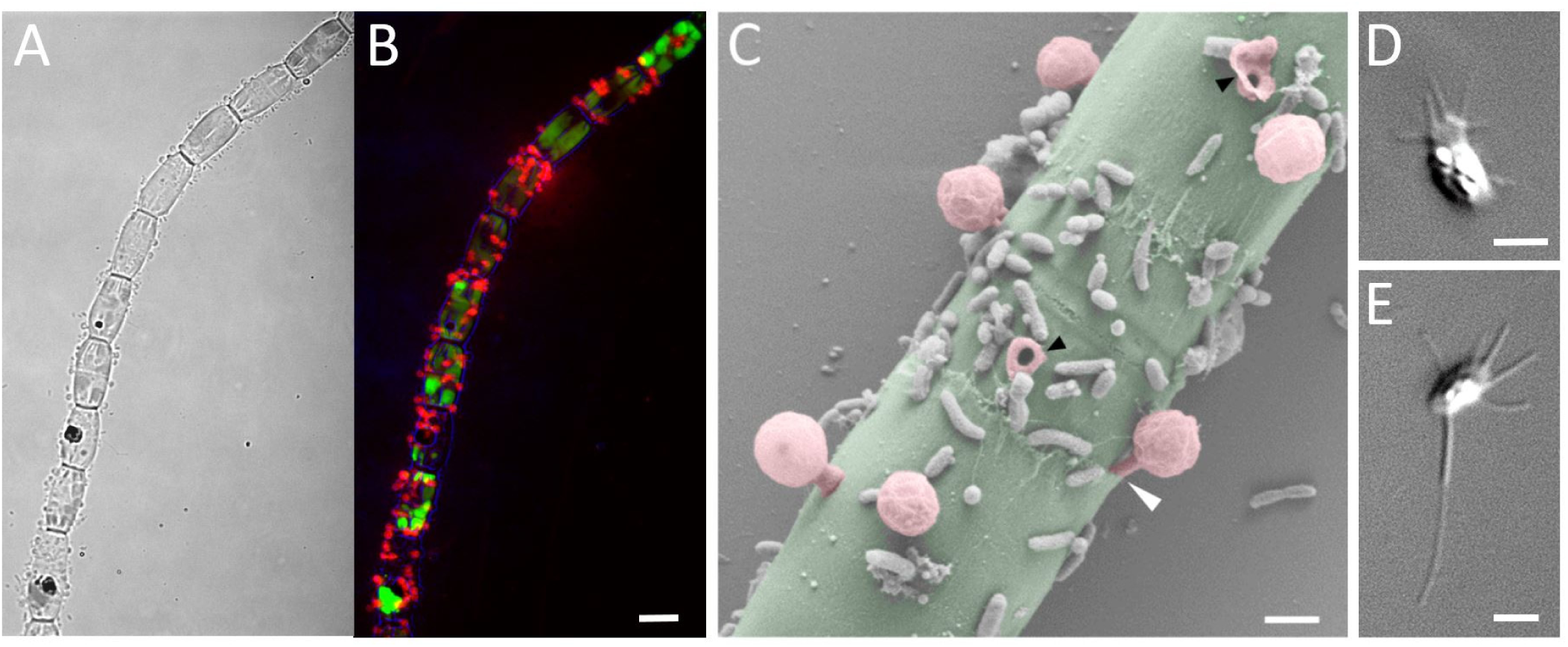
Zoospores and chitin-bearing infective cysts of *Paraphelidium tribonemae*. (A) Filament of *Tribonema gayanum* infected by *P. tribonemae* seen under optical microscopy and (B) the same filament stained with fluorescent wheat germ agglutinin (WGA) showing the presence of chitin in infective cysts under epifluorescence microscopy. (C) False-colored scanning-electron microscopy image of a filament infected by several *P. tribonemae* cysts (pedunculated rounded structures). The algal host filament is colored in green and parasite cysts in pink. Note that one cyst germ tube is penetrating the host cell by a cell-wall juncture (white arrowhead) and two cysts are broken (black arrowheads), showing the penetration channel. (D) Amoeboid zoospore (infrequent). (E) Amoeboflagellated zoospore. Scale bar: A = 5 µm, B-F = 1 µm. Phylogenetic trees related to chitin and other cell-wall synthesis and degradation-related enzymes are shown in Supplementary Fig. 2. Zoospore motility can be seen in Supplementary Video1.

Among the rest of chitin-related enzymes, we identified twelve sequences (at least five different homologs) of chitin deacetylase^25,26^ (Supplementary Fig. 2b). We detected at least three class II chitinase homologs (eight total sequences), which are ancestral in opisthokonts, containing the Glyco_hydro_18 (PF00704) domain, and a class I chitinase (CTSI) with a Glyco_hydro_19 (PF00182) domain^27^. The latter included the catalytic site, an N-terminal predicted signal peptide and a transmembrane region, suggesting an extracellular chitinase activity. CTSI has a peculiar phylogenetic distribution in eukaryotes, occurring only in Viridiplantae, Fungi, Opisthosporidia and Ecdysozoa (Supplementary Fig. 2c). *Rozella* contains two homologs and Microsporidia at least one; they have N-terminal signal peptides and are predicted to localize extracellularly but lack transmembrane domains. In our phylogenetic tree, opisthosporidian sequences appeared scattered within metazoan and fungal sequences. This might be the result of hidden paralogy and/or horizontal gene transfer (HGT) (Supplementary Fig. 2c). Regardless its evolutionary origin, aphelid CTSI might be involved in the self-degradation of resting spore and cyst wall chitin. This might happen both, during their release from chitin-containing resting spores or at the tip of the germ tube during infection, as previously suggested for *Rozella*^9^ (Fig. 1).

Although not found in chytrids, 1,3-beta-glucan is also considered an idiosyncratic fungal cell-wall component^19^. Surprisingly, we identified two 1,3-beta-glucan synthase (FKS1) homologs (four sequences), with split Glucan_synthase (PF02364) and FKS1_dom1 (PF14288) domains (fused for the phylogenetic analysis) (Supplementary Fig. 2d). The presence of FKS1, absent in *Rozella* and Microsporidia, in aphelids traces its origin back to the ancestor of Fungi and aphelids.

To feed on the algal cytoplasm, aphelids need to traverse the algal cell wall but, so far, the specific penetration mechanism, whether mechanical (germ-tube penetration at algal cell-wall junctures) or enzymatic (digestion of algal cell-wall components) was uncertain^1,2^. Scanning electron microscopy (SEM) observations showed both, clear cases of mechanical penetration via junctures but also infecting cysts scattered on the algal cell-wall surface far from junctures (Fig. 2c). SEM images additionally confirmed WGA-epifluorescence observations of multiple parasitoid cysts co-infecting the same host cell (Fig. 2a-c). Multiple infections open the intriguing possibility for aphelids to have sex within the algal host. *Tribonema* cell walls contain cellulose II based on 1,6-linked glucan (alkali soluble cellulose), 1,3 and 1,4-linked xylose, 1,3-linked rhamnose and, mostly, 1,3, 1,4 and 1,6-linked glucose^28^. We performed sequence similarity searches of known fungal cellulases^29^ using the database mycoCLAP^30^, which contains functionally characterized proteins, to look for these enzymes in aphelids, followed by phylogenetic analyses. In support of an enzymatic algal cell-wall penetration, we identified various cellulases in the *Paraphelidium* transcriptome belonging to glycoside-hydrolase families GH3 and GH5. We detected three homologs of the GH3 cellulase beta-glucosidase/xylosidase^31^, which is not found in other opisthosporidians but is present in fungi, amoebozoans, several opisthokonts and other protists, as well as bacteria. Our phylogenetic analysis shows that the three aphelid sequences are most closely related to deep-branching opisthokont protists (respectively, *Capsaspora*, choanoflagellates, nucleariids) (Supplementary Fig. 2e). Additionally, we identified at least three GH5 cellulase^32^ homologs (seven sequences in total) in *P. tribonemae*, which were related to GH5 widespread in fungi (GH5_11, GH5_12 and GH5_24) (Supplementary Fig. 2f). Collectively, these observations strongly suggest that these cellulases are involved in the alga cell-wall penetration, but direct proof will only be obtained by purifying or heterologously expressing those cellulases and testing their activity in vitro.

In summary, *Paraphelidium* has a complex life cycle with chitin-containing infective cysts and a diversified set of enzymes involved in chitin metabolism and possibly cellulose hydrolysis.

### Primary metabolism reminiscent of free-living lifestyles

Analysis of the *Rozella allomycis* genome showed that, like microsporidian parasites, it has significantly reduced metabolic capabilities^9^. To comparatively assess the metabolic potential of aphelids, we investigated the presence of orthologous groups (OGs) related to eight primary metabolism categories (Gene Ontology) in the transcriptome of *Paraphelidium*, using eggNOG annotation^33^. We thus identified 1,172 OGs in *Paraphelidium* and a set of 41 eukaryotic species including representatives of major fungal lineages, opisthokont protists and other eukaryotic parasites (Supplementary Table 2). Based on their OG distribution, we built a dissimilarity matrix that was analyzed by Principal Coordinate Analysis. The first axis clearly separated *Paraphelidium* from Microsporidia, *Mitosporidium*, *Paramicrosporidium* and *Rozella*, the latter two positioned near one another and having an intermediate position similar to other protist parasites (e.g. *Trypanosoma, Leishmania, Toxoplasma*) (Fig. 3a and Supplementary Fig. 3a). *Paraphelidium* positioned at the same level as fungi, *Capsaspora, Corallochytrium* and *Parvularia*, along axis 1. However, axis 2 separated *Paraphelidium* and fungi from the rest of eukaryotes. These relationships were further visualized in cluster analysis of pairwise species comparisons (Supplementary Fig. 3b). The PCoA suggested that *Paraphelidium* had a rich metabolic gene complement, which was made evident by the OG presence/absence heatmap showing that aphelids have a metabolic potential fully comparable to that of (especially chytrid) fungi (Fig. 3b).

**Figure 3.**
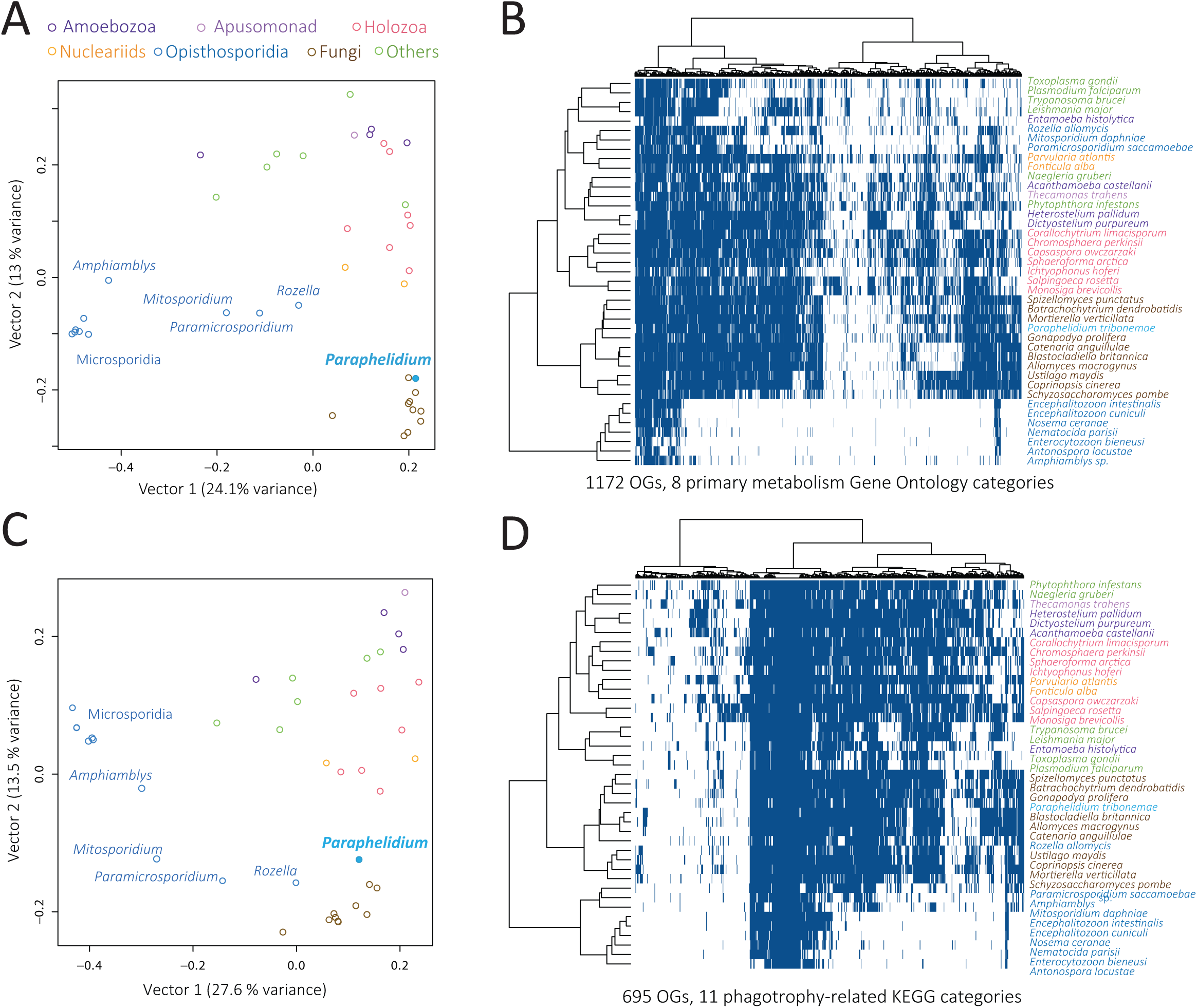
Complexity of *Paraphelidum tribonemae* metabolism and cytoskeleton-trafficking-phagotrophy-related proteome. (A) Principal Coordinate Analysis (PCoA) and (B) binary heat-map and species clustering based on the presence/absence of 1172 orthologous genes (OGs) belonging to 8 primary metabolism Gene Onthology categories across 41 eukaryotic genomes/transcriptomes. (C) PCoA and (D) binary heat-map and species clustering based on the presence/absence of 695 KEGG orthologs (OGs) related to cytoskeleton, membrane-trafficking and phagotrophy, which were selected from 11 KEGG categories. Species are color-coded according to their taxonomic assignment.

The most distinctive categories when comparing *Paraphelidium* and other Opisthosporidia were “energy production and conversion” followed by “amino acid, nucleotide and lipid transport and metabolism”. In all metabolic categories, the aphelid clustered with fungi, and more specifically with chytrids, and sometimes with other free-living opisthokonts (e.g. nucleariids, *Capsaspora*). By contrast, *Rozella* always clustered with *Mitosporidium* and *Paramicrosporidium* either together with other Microsporidia or with other parasitic protists (Supplementary Fig. 3c-f). The only exception corresponded to *Paramicrosporidium* which, for “energy production and conversion”, clustered with nucleariids (Supplementary Fig. 3c), in agreement with their rich energy-related gene set^17^.

To check whether these differences between *Paraphelidium* and *Rozella* affected particular metabolic pathways, we compared the annotated proteins in the two organisms based on KEGG annotation^34^. The comparison of the KEGG general metabolic pathway map showed that, even accounting for the possibility that we missed some genes in the *Paraphelidium*’s transcriptome (e.g. mitochondrial-encoded proteins), the aphelid map contained 200 OGs more than *Rozella* (548 vs 348 OGs) (Supplementary Fig. 3g). In agreement with previous observations, major differences were observed in “energy production and conversion”, and “amino acid, nucleotide and lipid transport and metabolism”. In particular, contrary to *Rozella*, which lacks most subunits of the mitochondrial electron transport chain complex I (NADH dehydrogenase; ETC-I)^9^, *Paraphelidium* possesses a practically complete ETC-I as inferred from the nuclear encoded transcripts (the *P. tribonemae* transcriptome is biased against mitochondrial transcripts, which lack polyA) (Supplementary Fig. 3h). *Paraphelidium* also possesses wide capabilities related to nucleotide (e.g. purine, uridine and inosine biosynthesis) and amino acid (serine, threonine, methionine, lysine, ornithine, histidine, shikimate pathway) metabolism, which *Rozella* has lost (Supplementary Fig. 3i). Likewise, the aphelid has also retained pathways for phosphatidylcholine, cholesterol and fatty acid biosynthesis that were subsequently lost in *Rozella*^9^. Most carbohydrate metabolic pathways were conserved in the two opisthosporidians, except for the galactose synthesis and degradation, also lost in *Rozella* (Supplementary Table 3).

By contrast, compared to *Rozella*, and under the assumption that our transcriptome is rather complete, the aphelid seems to lack several enzymes involved in catecholamine biosynthesis (Supplementary Table 3). However, some of these are also absent from *Capsaspora, Monosiga, Salpingoeca* or *Spizellomyces*. These compounds are likely involved in cell-cell communication in microbes^35^, e.g. parasite-host signaling, suggesting that they might have a role in rozellid parasitism. The aphelid seems to lack other parasite-specific proteins, such as crinkler, nucleoside H^+^-symporters or ATP/ADP-antiporters, which occur in *Rozella* and/or Microsporidia^9^.

These observations suggest that *Paraphelidium* has a complex metabolic profile, being functionally closer to free-living protists than to parasites and having affinities with fungi and, to a lesser extent, nucleariids and holozoan protists.

### Distinct and ancestral-like phagotrophy-related machinery

Like rozellids, aphelids are phagotrophs, but their global metabolism resembles that of osmotrophic fungi. What does their phagocytosis-related proteome look like? The core phagocytic molecular machinery, already present in the last eukaryotic common ancestor^36^, is involved in multiple dynamic cell processes (endocytosis, exocytosis, autophagy, protein recycling, etc.). Structurally, the phagocytic machinery encompasses various endomembrane organelles (phagosomes, lysosomes, peroxisomes, endoplasmic reticulum, Golgi apparatus), and multiple membrane-trafficking components (signaling pathways, transporters, specific enzymes, cytoskeleton elements, motors, etc.)^37^. To look for phagotrophy-related genes in *Paraphelidium* and the 41 additional eukaryotes used for comparison, we built a presence/absence matrix of 695 KEGG orthologs (OGs) from 11 functional categories (5 KEGG maps and 6 BRITE categories) that aimed at including all necessary proteins to perform phagotrophy; i.e. phagolysosome biogenesis, membrane trafficking and the actin cytoskeleton^37^ (Supplementary Table 4). A PCoA showed that *Paraphelidium* and *Rozella* are much closer to one another in terms of phagotrophy-than metabolism-related genes, grouping with fungi and far from Microsporidia (Fig. 3c and Supplementary Fig. 4a). This pattern was also evident in the presence/absence matrix (Fig. 3d and Supplementary Fig. 4b). In both, the two opisthosporidians clustered with early-branching fungi: chytridiomycetes (*Spizellomyces*, *Batrachochytrium*, *Gonapodya*) and blastocladiomycetes (*Allomyces*, *Catenaria*, *Blastocladiella*). Overall, *Paraphelidium* and *Rozella* have similar, but not identical, protein sets involved in phagosome, lysosome and endocytic processes. This suggests that, from a common phagocytic ancestor, different protein subsets specialized in both lineages for the phagocytosis function (Supplementary Fig. 4c-e).

**Figure 4.**
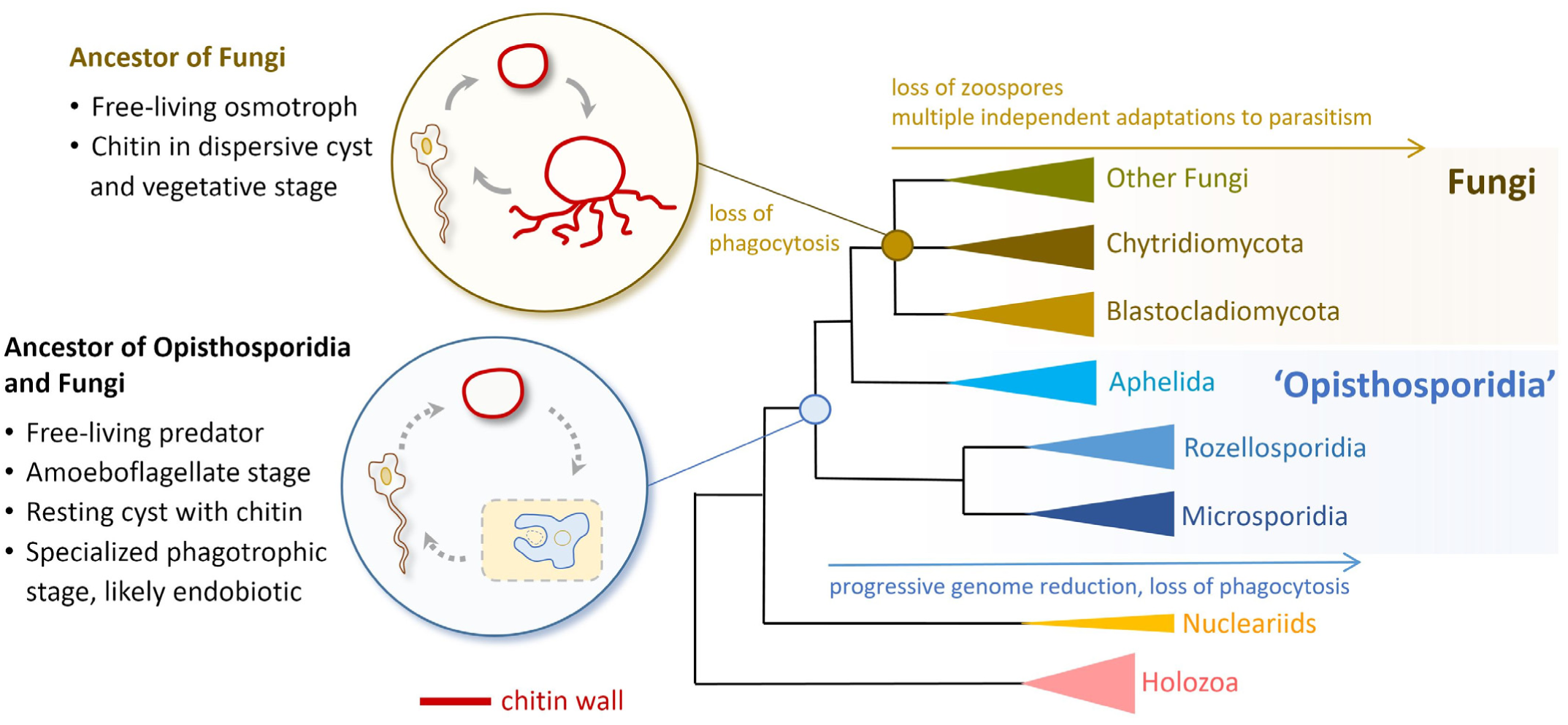
Early evolution of Fungi and related lineages. Schematic representation of evolutionary relationships between Fungi, rozellids and Microsporidia, within the holomycotan branch of Opisthokonts. Inferred key ancestral features and life cycle stages are depicted at the corresponding ancestral nodes.

In order to gain more insights into the diversification of the actin cytoskeleton toolkit of fungi and opisthosporidians, we analyzed the evolution of myosin motor proteins. The myosin toolkit of *Paraphelidium* contains a mixture of classical fungal families and others previously identified in holozoans (animals and their unicellular relatives; Supplementary Table 5). We recovered diversified class I myosins in *Paraphelidium*, *Spizellomyces* and nucleariids (Supplementary Fig. 4g), with paralogs of Ic/h and Ik families, previously described only in holozoans^38^. *Paraphelidium,* nucleariids and *Gonapodya* also possess homologs of class XV myosins, formerly thought holozoan-specific^38^. In addition, the aphelid not only possesses homologs of the V/VII myosin-motor family associated to chitin-synthase (see above; Supplementary Fig. 2a), but also myosins of If (pan-eukaryotic), II (amorphean), and XXII (opisthokont) classes, which clustered with previously described fungal homologs (Supplementary Table 5; Supplementary Fig. 4f). Thus, compared with the ancestral opisthokont myosin complement^38^, *Paraphelidium* retains all but one myosin class (VI), with homologs of all myosin families present in fungi (If, II, V, XVII - chitin synthase) plus four additional families (Ic/h, Ik, XV and XXII) that were lost in “eumycotan” fungi (i.e., fungi to the exclusion of chytrids). This suggests an independent step-wise simplification of the myosin complement in fungi, *Rozella* and Microsporidia, with *Paraphelidium,* nucleariids and chytrids retaining ancestral classes.

Recently, it has been proposed that WASP and SCAR/WAVE genes, encoding activators of branched actin assembly proteins^39^, are essential to build actin-based pseudopodia (filopodia), being fundamental for the typical movement of filopodiated cells called “alpha-motility”^40^ (Supplementary Table 1). The aphelid zoospore motility (Supplementary Video 1) resembles that of chytrid fungi^40^ and its transcriptome contains homologs of WASP and SCAR/WAVE. Interestingly, although *Rozella* also contains WASP and SCAR/WAVE homologs, filopodia have not been described in its zoospores^41–43^. If *Rozella* truly lacks filopodia, this might call into question the proposed essential role of these genes for filopodial movement^40^ and might instead suggest their potential involvement in phagotrophy in this organism.

### Aphelids and the free-living-like ancestor of Fungi

From an evolutionary perspective, the current situation in the holomycotan branch (including Fungi) of the eukaryotic super-group Opisthokonta mirrors that of the Holozoa (including Metazoa), where the discovery of deeply-branching unicellular protists that possess genes thought unique to animals continues to challenge previous evolutionary schemes about the emergence of Metazoa^15,44^. Thus, the discovery that widely diverse environmental sequences formed a deeply-branching clade that appeared sister to fungi and that included theparasite *Rozella allomyces* (named rozellids^3^, and subsequently Cryptomycota^4^, Rozellomycota^45^ or Rozellosporidia^46^), triggered the discussion of what Fungi actually are^19,47^. A debate further nourished by the discovery that aphelids, another widely diverse group of parasites of algae^48^, formed a large clade with rozellids and Microsporidia based on rRNA and RNA-polymerase genes^1,5,46,49^. This seemingly monophyletic clade was named Opisthosporidia and branched as sister to classical Fungi (the monophyletic clade including from chytrids and their relatives to the Dikarya^47^) in phylogenetic trees^1^. Lately, many mycologists include the three opisthosporidian lineages, Aphelida, Rozellosporidia and Microsporidia, within Fungi^9,19,50,51^. Some authors even incorporate as fungi the free-living phagotrophic chitin-lacking nucleariids, thus pushing the limits of Fungi to incorporate all holomycotan (possibly a confusing name) lineages, despite asserting that “…the kingdom Fungi is characterized by osmotrophic nutrition across a chitinous cell wall…”^52^. However, unlike fungi, aphelids and rozellids (the deepest branches in the holomycotan clade to the exclusion of nucleariids) are phagotrophs, lack a chitin cell wall at the vegetative stage and are endobiotic, having a unique mode of penetration into the host^1,2,47,49^. Also, because *Rozella* has a very reduced genome, a typical trait of streamlined parasites, some authors inferred a parasitic nature for the ancestor of Fungi^9^.

The study of the *Paraphelidium tribonemae* transcriptome adds some light to this controversy. Our multi-gene phylogenetic analyses place the aphelids as a pivotal group branching very deeply in the Opisthosporidia/Fungi clade. However, contrary to previous phylogenetic analyses based on few genes, Opisthosporidia do not seem monophyletic. Aphelids strongly emerge as the sister group to Fungi to the exclusion of rozellids and Microsporidia (Fig. 1). Our results suggest that Fungi evolved from ancestors that were related to, or potentially were, aphelids. Because of its deep pivotal position, the comparative study of the *Paraphelidium* transcriptome allows better inferring ancestral states for Fungi and the larger Opisthosporidia clade (Fig. 4). *Paraphelidium* has a complex metabolism resembling that of free-living chytrids but feeds by phagotrophy as free-living nucleariids and holozoan protists (Fig. 3). This suggests that aphelids are ‘borderline’ parasites (or parasitoids, according to some definitions) that have not undergone the reduction process that characterizes *Rozella* and all members along the microsporidian branch^6,8,9^. Being more gene-rich and close to the root, aphelids may have retained more ancestral features. This advocates for a free-living opisthosporidian ancestor that had a complex life cycle including chitin-containing resting cysts, amoeboflagellate zoospores and a phagotrophic amoeba stage possibly specialized in endobiotic predation (Fig. 4). By contrast, the fungal ancestor was a free-living osmotroph that had amoeboflagellate zoospores and chitin in the vegetative stage. From their aphelid-like opisthosporidian free-living ancestor, the fungal lineage lost phagotrophy, acquiring its ecologically successful osmotrophy whereas, on the rozellid line, endobiotic phagotrophic predation shifted into obligate parasitism with more complex life cycles, highly specialized morphologies and genome and metabolic reduction. The analysis of additional genomes/transcriptomes for other aphelids, rozellids and deep-branching fungi should help establishing a solid phylogenomic framework to validate and refine this evolutionary scenario

## Methods

### Cultures

*Paraphelidium tribonemae* was maintained in an enriched culture with its host *Tribonema gayanum* in mineral medium or in Volvic™ at room temperature in the presence of white light^2^. The culture was progressively cleaned from other heterotrophic eukaryotes by micromanipulation and transfer of infected host filaments to uninfected *T. gayanum* cultures.

### WGA staining, epifluorescence and scanning electron microscopy

To detect chitin in different cell cycle stages of *Paraphelidium tribonemae,* we incubated actively growing cultures with 5µg/mL Wheat Germ Agglutinin (WGA) conjugated to Texas Red (Life Technologies) for 10 minutes at room temperature. After rinsing with Volvic™ water, cells were then observed under a LEICA DM2000LED fluorescence microscope with an HCX_PL_FLUOTAR 100x/1.30 oil PH3 objective. Pictures were taken with a LEICA DFC3000G camera using the LEICA Application Suite v4.5 and edited with ImageJ (http://imagej.nih.gov/ij/). For scanning electron microscopy (SEM) observations, we transferred 1-week-old cultures to a Petri dish containing mineral medium and two coverslips. Cells were let to settle overnight before fixation (1% OsO_4_ and 1% HgCl_2_ for 45 min). After washing (3×10 min in distilled water), samples were dehydrated in ethanol series (30%, 50%, 70%, 90%, 96%, 100%) for 10 minutes each. After critical-point drying and sputter-coating with platinum, cells were visualized with a Zeiss Sigma FE-SEM at 1kV acceleration voltage.

### Transcriptome Sequencing

To obtain a complete representation of the aphelid transcriptome across different cell-cycle stages, we extracted RNA from a young culture (3 days after inoculation) rich in aphelid zoospores and cysts, and eight old cultures (5-7 days after inoculation) with few remaining *T. gayanum living* cells and rich in aphelid plasmodia, resting spores and zoospores. To minimize algal overrepresentation, we maintained the cultures in the dark. Total RNA was extracted with the RNeasy mini Kit (Qiagen), quantified by Qubit (ThermoFisher Scientific) and sent to Eurofins Genomics (Germany) for *de novo* transcriptome sequencing. Two cDNA Illumina libraries for, respectively, the ‘young’ and ‘old’ enrichment cultures were constructed after polyA mRNA selection and paired-end (2×125 bp) sequenced with Illumina HiSeq 2500 Chemistry v4.

### Transcriptome assembly, decontamination and annotation

A total of 71,015,565 and 58,826,379 reads, respectively, were obtained for the libraries of ‘young’ and ‘old’ cultures. From these, we obtained 34,844,871 (‘young’) and 25,569,845 (‘old’) paired-end reads. After quality/Illumina Hiseq adapter trimming using Trimmomatic v0.33^53^, we retained 8,601,196 forward and 6,535,164 reverse reads (‘young’) and 10,063,604 forward and 4,563,537 reverse unpaired reads (‘old’). Resulting reads were assembled using Trinity^54^ with “min_kmer_cov 2” and “normalize_reads” options. This initial assembly, Partr_v0, contained 68,130 contigs. 47 of them were identified as ribosomal RNAs using RNAMMER^55^ and included sequences of *Paraphelidium tribonemae*, its host *Tribonema gayanum*, chloroplasts and bacteria. No other eukaryotic ribosomal genes were detected. To decontaminate *P. tribonemae* from host and bacterial sequences, we used Blobtools^56^ and, as input files, our assembly, a read map file obtained with the fast gapped-read alignment program Bowtie2^57^ and a formatted hit list obtained after applying diamond-blastx^58^ to our sequences against the NCBI RefSeq database (e-value threshold 1e-10). To build the blobDB file, we used the first 10 hits for each putative transcript with the bestsum as tax-rule criterion to get a taxonomic affiliation at species rank. Sequences with no hit (41,836 sequences, 61.4% of the assembly) and sequences affiliated to bacteria, archaea or viruses were removed. The remaining 21,688 eukaryotic sequences were blasted against a partial REFSEQ protein database (e-value threshold 1e-10) containing sequences of 5 fungi (*Spizellomyces punctatus* DAOM BR117, *Batrachochytrium dendrobatidis* JAM81, *Agaricus bisporus* var. *bisporus* H97, *Cryptococcus neoformans* var. *grubii* H99 and *Cryptococcus neoformans* var. *neoformans* JEC21) and 6 stramenopiles (*Nannochloropsis gaditana* CCMP526, *Phytophthora sojae, Phytophthora parasitica* INRA-310, *Phytophthora infestans* T30-4, *Aureococcus anophagefferens* and *Aphanomyces invadans*). If the first hit was a stramenopile (potential host origin), the sequence was discarded. 13,786 transcripts appeared free of contamination. Transdecoder v2 (http://transdecoder.github.io) with --search_pfam option yielded 16,841 peptide sequences that were filtered using Cd-hit v4.6^59^ with a 100% identity resulting in 13,035 peptides. These peptides were functionally annotated with eggNOG-mapper^33^ using both DIAMOND and HMMER mapping mode, eukaryotes as taxonomic scope, all orthologs and non-electronic terms for Gene Ontology evidence. This resulted in 10,669 annotated peptides for the *Paraphelidium tribonemae* predicted proteome (version Partr_v1). Finally, we generated transcriptomic data for the *Tribonema gayanum* host using the same methods and parameters for sequencing, assembly and annotation, as described above. After blasting Partr_v1 against a database of *Tribonema gayanum* proteins, we further excluded 219+11 additional proteins that were 100% or >95% identical to *Tribonema* proteins. After these additional cleaning steps, version 1.5 of the *Paraphelidium tribonemae* predicted proteome contained 10,439 proteins (Supplementary Table 1). The completeness of this new transcriptome was estimated to 91.4% with BUSCO^10^ using the eukaryotic ortholog database 9 (protein option).

### Phylogenomic analyses

We used three different protein datasets previously used for phylogenomic analyses of Opisthokonta and, more specifically, Microsporidia that included, respectively, 93 single-copy protein domains (SCPD)^11^, 53 proteins (BMC)^18^ and 259 proteins (GBE)^8^. We updated these datasets with sequences from the *Paraphelidium tribonemae* transcriptome, thirteen derived Microsporidia as well as *Mitosporidium daphniae*^12^, *Paramicrosporidium saccamoebae*^17^and *Amphiamblys* sp.^8^. Some stramenopile taxa such *Ectocarpus siliculosus*^60^, *Thalassiosira pseudonana*^61^ or *Phytophthora infestans* (GenBank NW_003303758.1) were also included to further prevent any previously undetected *Tribonema gayanum* contamination. Markers were aligned with MAFFT v7^62^, using the method L-INS-i with 1000 iterations. Unambiguously aligned regions were trimmed with TrimAl^63^ with the automated1 algorithm, then approximate Maximum Likelihood (ML) trees were inferred using FastTree^64^, all visually inspected in Geneious^65^ to check for possible contamination. Contamination-free alignments were relieved from distant outgroup taxa. For in-depth analyses, we used opisthokont, mainly holomycotan, sequences, including apusomonad and breviate sequences as outgroup. We then made phylogenomic analyses including or excluding thirteen fast-evolving microsporidian parasites (49 or 36 species). Sequences of the selected taxa were concatenated using Alvert.py from the package Barrel-o-Monkeys (http://rogerlab.biochemistryandmolecularbiology.dal.ca/Software/Software.htm#Monkeybarrel). This resulted in alignments containing the following number of amino acids as a function of dataset and taxon sampling: 22,976 (SCPD49); 15,744 (BMC49); 87,045 (GBE49); 24,413 (SCPD36); 16,840 (BMC36); 96,029 (GBE36). Bayesian phylogenetic trees were inferred using PhyloBayes-MPI v1.5^66^ under the CAT-Poisson evolutionary model, applying the Dirichlet process and removing constant sites. Two independent MCMC chains for each dataset were run for >15,000 generations, saving one every 10 trees. Phylogenetic analyses were stopped once convergence thresholds were reached after a burn-in of 25% (i.e., maximum discrepancy < 0.1 and minimum effective size > 100 using bpcomp). We also applied ML phylogenetic reconstruction using IQ-TREE^67^ with the profile mixture model C60^14^. Statistical support was obtained with 1,000 ultrafast bootstraps^16^ and 1000 replicates of the SH-·like approximate likelihood ratio test^68^. To alleviate the local computational burden, many of these analyses were carried out using CIPRES Science Gateway^69^. Trees were visualized with FigTree (http://tree.bio.ed.ac.uk/software/figtree/). To test if the best topology obtained was significantly better than other alternatives, we tested whether constrained alternative topologies could be rejected for different datasets. We used Mesquite^70^ to constrain topologies representing the opisthosporidian monophyly (aphelids+rozellids+microsporidians) and the monophyly of aphelids with either blastocladiomycetes, chytridiomycetes or chytridiomycetes+blastocladiomycetes. The constrained topologies without any branch-length were reanalyzed with the “-g” option of IQ-TREE and the best-fitting model for each dataset. The resulting trees were then concatenated and AU tests were performed for each dataset with the “-z” and “-au” options as described in the advanced documentation of IQ-TREE. To minimize systematic error due to the inclusion of fast-evolving sites in our protein datasets, we progressively removed the fastest evolving sites at steps of 5% sites removed at a time. Among-site evolutionary rates were inferred using IQ-TREE “-wsr” option and its best-fitting model for each dataset (Supplementary Table 2). A total of 19 subsets were created for each dataset. We then reconstructed phylogenetic trees using IQ-TREE with the same best fitting model as for the whole dataset. To know how supported were alternative topologies in bootstrapped trees, we used CONSENSE from the PHYLIP package^71^ and interrogated the .UFBOOT file using a Python script (M Kolisko, pers. comm).

### Homology searches and phylogenetic analyses of specific proteins

Protein sets used in this study were obtained in May 2017 from the NCBI protein database (https://www.ncbi.nlm.nih.gov/protein/), except the following: *Chromosphaera perkinsii*, *Corallochytrium limacisporum* and *Ichthyophonus hoferi* genome data, obtained from^15^; *Spizellomyces punctatus, Gonapodya prolifera, Batrachochytrium dendrobatidis, Allomyces macrogynus, Catenaria anguillulae* and *Blastocladiella britannica*, retrieved from MycoCosm portal, Joint Genome Institute* (http://www.jgi.doe.gov/); *Parvularia atlantis* (previously *Nuclearia* sp. ATCC50694, from https://doi.org/10.6084/m9.figshare.3898485.v4); and *Paramicrosporidium saccamoebae,* from NCBI in January 2018. [* These sequence data were produced by the US Department of Energy Joint Genome Institute http://www.jgi.doe.gov/ in collaboration with the user community]. Query sequences to retrieve chitin and related cell-wall proteins (chitin synthases – CHS –, chitin deacetylases – CDA – and chitinases – CTS –) were obtained from the study of *Rozella allomycis*^9^ except for 1,3-beta glucan synthase (FKS1), which was obtained from *Saccharomyces cerevisiae* S288C. We used sequences from *Aspergillus fumigatus* as initial queries for cellulases^29^ and also opisthokont homologs retrieved by BLASTp against the *Paraphelidium* transcriptome. Homology was confirmed by searching for the identified sequences in the eggNOG^33^ annotation file (Supplementary Table 1), performing reverse BLAST against the non-redundant NCBI database and HMMER analyses^72^. Cell-wall proteins, notably chitinases and cellulases, were also annotated using CAZyme identification tools (http://csbl.bmb.uga.edu/dbCAN/annotate.php). We produced comprehensive alignments including sequences retrieved from previous studies and sequences identified by BLAST in GenBank or in the mycoCLAP database^30^ with the MAFFT web server^62^ applying the progressive L-INS-i algorithm (except for the multidomain CHS sequences for which E-INS-i was used). Alignments were trimmed from gaps and ambiguously aligned sites using Trimal^63^ with the automated1 algorithm. ML trees were inferred using IQ-TREE web server^67^ with the LG+G4 evolutionary model and visualized with FigTree. In the case of myosins, protein sequences were retrieved by querying the HMM profiles of the myosin head domain (Pfam accession: PF00063) against our database of *Paraphelidium tribonemae* proteins, using the *hmmersearch* utility of HMMER (http://hmmer.org) and the profile-specific gathering threshold cut-off. For each retrieved homolog, the protein segment spanning the Pfam-defined myosin head domain was aligned against a pan-eukaryotic alignment of myosins^38^, using MAFFT^62^ with the G-INS-i algorithm optimized for global sequence homology. The alignment was run for up to 10^3^ cycles of iterative refinement and manually examined, curated and trimmed. The ML phylogenetic reconstruction of the trimmed alignment (997 sequences, 400 sites) was done with IQ-tree^67^ under the LG model with a 10-categories free-rate distribution, selected as the best-fitting model according to the IQ-TREE *TESTNEW* algorithm as per the Bayesian information criterion. The best-scoring tree was searched for up to 100 iterations, starting from 100 initial parsimonious trees; statistical supports for the bipartitions were drawn from 1000 ultra-fast bootstrap^16^ replicates with a 0.99 minimum correlation as convergence criterion, and 1000 replicates of the SH-like approximate likelihood ratio test. We classified the myosin homologs of *Paraphelidium tribonemae* according to the phylogenetic framework of Sebé-Pedrós et al.^38^. To characterize the classified homologs (Supplementary Table 5), we recorded the protein domain architectures of the full-length proteins using Pfamscan and the 29th release of the Pfam database^73^.

### Comparative analyses of proteins involved in primary metabolism, cytoskeleton, membrane-trafficking and phagotrophy

To get a broad view of the metabolic capabilities of *Paraphelidium tribonemae* in comparison with parasitic and free-living organisms, we performed a statistical multivariate analysis similar to that previously used for a similar purpose in *Rozella allomycis*^9^. We initially searched in *P. tribonemae* for the presence of 6,469 unique orthologous groups as per eggNOG^33^, corresponding to 8 primary metabolism categories (Gene Ontology, GO). The correspondence between GO terms and primary metabolism COGs was obtained from http://geneontology.org/external2go/cog2go, and was as follows: [C] Energy production and conversion (912 OGs); [G] Carbohydrate transport and metabolism (1901 OGs); [E] Amino acid transport and metabolism (714 OGs); [F] Nucleotide transport and metabolism (351 OGs); [H] Coenzyme transport and metabolism (375 OGs); [I] Lipid transport and metabolism (965 OGs); [P] Inorganic ion transport and metabolism (842 OGs) and [Q] Secondary metabolites biosynthesis, transport and catabolism (570 OGs). From these categories, we identified 1,172 non-ambiguous OGs in the *P. tribonemae* transcriptome that were shared with a set of other 41 protist species (Supplementary Table 3). These 41 protist species represent a broader taxonomic sampling as compared to the *R. allomyces* study^9^. We included 11 opisthosporidians: *Paramicrosporidium saccamoebae* (Par_sa), *Amphiamblys* sp. (Amp_sp), *Antonospora locustae* (Ant_lo), *Encephalitozoon cuniculi* (Enc_cu), *Encephalitozoon intestinalis* (Enc_in), *Enterocytozoon bieneusi* (Ent_bi), *Mitosporidium daphniae* (Mit_da), *Nematocida parisii* (Nem_pa), *Nosema ceranae* (Nos_ce), *Rozella allomycis* (Roz_al), *Paraphelidium tribonemae* (Par_tr); 10 fungi covering most fungal phyla: *Spizellomyces punctatus* (Spi_pu), *Gonapodya prolifera* (Gon_pr), *Batrachochytrium dendrobatidis* (Bat_de), *Allomyces macrogynus* (All_ma), *Catenaria anguillulae* (Cat_an), *Blastocladiella britannica* (Bla_br), *Mortierella verticillata* (Mor_ve), *Coprinopsis cinerea* (Cop_ci), *Ustilago maydis* (Ust_ma), *Schizosaccharomyces pombe* (Sch_po); 2 nucleariids *Fonticula alba* (Fon_al), *Parvularia atlantis* (Par_at); 7 holozoans: *Salpingoeca rosetta* (Sal_ro), *Monosiga brevicollis* (Mon_br), *Capsaspora owczarzaki* (Cap_ow), *Chromosphaera perkinsii* (Chr_pe), *Corallochytrium limacisporum* (Cor_li), *Ichthyophonus hoferi* (Ich_ho), *Sphaeroforma arctica* (Sph_ar); 3 amoebozoans: *Acanthamoeba castellanii* (Aca_ca), *Dictyostelium purpureum* (Dic_pu), *Polysphondylium pallidum* (Pol_pa), *Entamoeba histolytica* (Ent_hi); the free-living apusomonad *Thecamonas trahens* (The_tr), *Naegleria gruberi* (Nae_gr) and 6 other eucaryotic parasites: *Toxoplasma gondii* (Tox_go), *Plasmodium falciparum* (Pla_fa), *Trypanosoma brucei* (Try_br), *Leishmania major* (Lei_ma), *Phytophora infestans* (Phy_in). We annotated all 41 protein sets using eggNOG-mapper^33^ using DIAMOND mapping mode, eukaryotes as taxonomic scope, all orthologs and non-electronic terms for Gene Ontology evidence. Species-wise OG counts were transformed into a presence/absence matrix (encoded as 0/1), or binary OG profile. This binary OG profile excluded all OGs with no identifiable orthologs in any of the 41 species used in this analysis (e.g., proteins related to exclusive prokaryotic metabolism or photosynthesis) leaving a total number of 1,172 primary metabolism OGs (Supplementary Table 3). The similarity between each species’ binary COG profile was assessed using the Pearson’s correlation coefficient (*r* statistic) as implemented in the R *stats* library^74^. We built a complementary species distance matrix by defining dissimilarity as 1-*r*., and we analyzed this dissimilarity matrix using a Principal Coordinate Analysis (PCoA) as implemented in the R *ape* library^75^, with default parameters. For each PCoA, we represented the two vectors with the highest fraction of explained variance, which were in all cases higher than the fractions expected under the broken stick model. In addition, we plotted binary OG profiles of each species in a presence/absence heatmap, produced using the heatmap.2 function in the R *gplots* library^76^. Species’ order was defined by a Ward hierarchical clustering of the aforementioned interspecific Pearson’s correlation coefficients. OGs’ order was defined by Ward hierarchical clustering on euclidean distances. All clustering and distance analyses were performed using R *stats* library^74^. We represented the raw species clustering (Pearson correlation+Ward) in a separate heatmap, scaling the color code to display positive Pearson correlation values (0 to 1). Finally, each sub-set of primary metabolism COGs was also analyzed separately (categories C, E, F, G, H, I, P and Q) using the same hierarchical clustering as above to cluster species according to their metabolism gene content (based on Pearson+Ward). We carried out similar comparisons and statistical analyses for proteins involved in phagotrophy, membrane trafficking and cytoskeleton. Briefly, we looked for proteins broadly related to those systems (actin cytoskeleton, exocytosis, fungal vacuole, exosome, endosome, mitochondrial biogenesis, ER-Golgi, lysosome, autophagy, peroxisome) in the KEGG BRITE reference database (http://www.genome.jp/kegg/brite.html). We identified 849 proteins related to membrane trafficking (ko04131), 279 cytoskeletal proteins (ko04812) and 361 proteins related to phagotrophy (KEGG map04144_endocytosis, map04145_phagosome, map04146_peroxisome, map04142_lysosome, map04810_actin_regulation, map04141_autophagy and map04071_sphingolipid_signal), which were used to carry out PCoA and clustering analyses as above (Supplementary Fig. 4). Since the eggNOG annotation file did not provide specific KOs (KEGG orthologs) but only KEGG maps, we used the following approach to get the KO profile for the 41 taxa. All proteins in 35 eukaryote genomes^77^ were clustered to a non-redundant set using cd-hit at a 90% identity threshold. Those 35 nr90 genomes were compared in an all-vs-all BLAST analysis with an e-value cutoff of 1×10^-3^. The all-vs-all BLAST output was clustered using the MCL algorithm with an inflation parameter of 2.0. To avoid lineage specific proteins, the resultant clusters were retained if the proteins in the cluster were derived from 3 or more organisms. Proteins from each cluster were aligned using MAFFT, the alignments trimmed with trimAl, and HMM profiles built for each alignment using HMMer3. This analysis resulted in a set of 14,095 HMMs. All proteins from the UniProt-SwissProt database were searched against the entire set of 14,095 HMMs. Each HMM was assigned a best-hit protein annotation from the UniProt-SwissProt database. KEGG ortholog (KO) identifiers (IDs) associated with each HMM were obtained from UniProtKB IDs and KO IDs using the mappings available from UniProtKB. Additional KO IDs were mapped to the HMMs by searching the HMMs against all proteins annotated to a given KO ID. The best hit bit score among those proteins to the HMM was compared to the best-hit bit score of the UniProtKB annotation for that HMM. If the best hit KO ID bit score was at least 80% of the best hit UniProtKB protein bit score, and there was no KO ID already associated with the UniProtKB annotation, the KO ID was transferred to that HMM. To build a presence-absence matrix associated with KO IDs, the HMMs were searched against all proteins in a genome. Best hit HMMs below threshold (total protein e-value <=1×10^-5^ and a best single domain e-value <=1e-4) were assigned for each protein in a genome. The best hit HMMs and their associated KO IDs were used for presence/absence calls. If at least one protein in a genome had a best hit to a given HMM, the associated KO was considered present for that genome. From a total of 1,568 OGs associated with the KEGG maps of interest, 633 KO IDs were identified in the UniProtKB annotations of the HMMs. An additional 62 KO IDs were mapped to HMMs by considering similarity of bit scores between the best UniProt/SwissProt protein hit to the HMM and the best KO ID protein hit to the HMM (Supplementary Table 3). Those 695 KO IDs were used to perform the same multivariate statistical analyses as per the metabolism section for the 41 taxa. The phagolysosome KEGG maps in *Paraphelidium tribonemae* and *Rozella allomycis* were compared using KEGG tools. To do so, we first annotated the two protein sets using BlastKOALA^78^ with eukaryotes as taxonomy group and genus_eukaryotes KEGG GENES database. We then uploaded our annotations in the KEGG Mapper Reconstruct Pathway and BRITE servers.

## Data availability

Raw read sequences have been deposited in NCBI under accession number PRJNA402032. *Paraphelidium tribonemae* metatranscriptomic nucleotide contig assembly and version 1.5 of the predicted proteome are deposited in figshare (https://figshare.com/authors/Guifr_Torruella/3846172) under Creative Commons 4.0 licence. All trees are available as supplementary file all_trees.txt

## Author contributions

G.T., D.M. and P.L-G. conceived and coordinated the study, and wrote the manuscript. G.T. performed culture cleaning, RNA extraction, *de novo* transcriptome assembly, phylogenomics and comparative genomic analyses. G.T. and X. G-B. cleaned the protein set from contamination. X. G-B. and A. S-B. performed myosin comparative genomic analyses. X. G-B. carried out multivariate statistical analyses of metabolic genes. G.T. and J.A.B. carried out phagolysosome protein analyses. G.T. and S.K. maintained cultures and performed WGA staining and imaging. S.K. studied morphology and life-cycle aspects, E.V. performed SEM fixation and imaging. All authors commented on the manuscript.

### Acknowledgements

We are grateful to Wayne Pfeiffer and Mark Miller from the CIPRES Science Gateway (http://www.phylo.org/) for computational resources and personal assistance. We thank Victoria S. Tcvetkova for micromanipulating infected *Tribonema* sp. filaments in view of establishing clean aphelid cultures, Alex de Mendoza for providing CHS alignments, Philippe Deschamps, Rafael Ponce, Guillaume Reboul, Ana Gutiérrez-Preciado, Ares Rocañín-Arjó and Ricardo Rodríguez de la Vega for discussion on bioinformatics and statistics aspects, and Peter Kubicek and Bernard Hernissat for helpful discussion about cellulases. Research leading to these results received funding from the European Research Council under the European Union’s Seventh Framework Program (ERC Grant Agreement 322669 ProtistWorld). GT was also financed by the European Marie Sklodowska-Curie Action (704566 AlgDates). S.K. was supported by RSF grant No 16-04-10302, ZIN RAS program AAAA-A17-117030310322-3. and “Jean d’Alembert” program (Université Paris-Saclay).

